# Modulation of MHC-E transport by viral decoy ligands is required for RhCMV/SIV vaccine efficacy

**DOI:** 10.1101/2020.09.30.321158

**Authors:** Marieke Verweij, Scott G. Hansen, Ravi Iyer, Nessy John, Daniel Malouli, David Morrow, Isabel Scholz, Jennie Womack, Shaheed Abdulhaqq, Roxanne M. Gilbride, Colette M. Hughes, Abigail B. Ventura, Julia C. Ford, Andrea N. Selseth, Kelli Oswald, Rebecca Shoemaker, Brian Berkemeier, William J. Bosche, Michael Hull, Jason Shao, Jonah B. Sacha, Michael K. Axthelm, Paul T. Edlefsen, Jeffrey D. Lifson, Louis J. Picker, Klaus Früh

**Author notes:** These authors contributed equally (Marieke Verweij, Scott G. Hansen, Ravi Iyer).

## Abstract

Strain 68-1 rhesus cytomegalovirus (RhCMV) vectors expressing simian immunodeficiency virus (SIV) antigens elicit CD8^+^ T cells that recognize peptide epitopes presented by major histocompatibility complex (MHC)-II and MHC-E molecules, instead of MHC-Ia, and are uniquely able to mediate stringent control and subsequent clearance of highly pathogenic SIV in ∼50% of vaccinated rhesus macaques (RMs). We show that the MHC-E ligand VMAPRTLLL (VL9), encoded by the Rh67 gene (or its HCMV UL40 counterpart) is required for recognition of RhCMV-infected fibroblasts by MHC-E-restricted CD8^+^ T cells via its ability to promote intracellular MHC-E transport. Moreover, deletion of Rh67 from 68-1 RhCMV/SIV vectors, or mutation of its embedded VL9 ligand, abrogated induction of MHC-E-restricted CD8^+^ T cell responses, leaving responses that exclusively target MHC-II-restricted epitopes. These MHC-II-presented CD8^+^ T cell responses, though comparable in response magnitude and functional differentiation to responses arising from the efficacious 68-1 vector, did not protect RMs against SIV challenge, indicating that Rh67/UL40-enabled direct priming of MHC-E-targeted CD8^+^ T cells is a crucial element of RhCMV/SIV vaccine efficacy.

**One Sentence Summary:** A cytomegalovirus protein (Rh67/UL40) that upregulates MHC-E expression on RhCMV/SIV-vector infected cells is required for induction of MHC-E-restricted CD8^+^ T cells and for protection against SIV.

---

Cytomegaloviruses (CMVs) are large β-herpesviruses that manifest extensive and complex immune evasion mechanisms to maintain viral persistence, while at the same time eliciting and maintaining robust anti-viral immune responses that restrain viral spread, the latter particularly including uniquely high frequency, circulating and tissue-based effector-memory T cell (T_EM_) responses (*1-5*). These robust T_EM_ responses can be harnessed against other pathogens or cancer by CMV-based vaccine vectors expressing heterologous antigens (*3, 6, 7*). Indeed, using strain 68-1 RhCMV/SIV vectors, we previously demonstrated an unprecedented type of SIV vaccine efficacy in which ∼50% of vaccinated RM exhibit early, complete replication arrest and subsequent clearance of highly pathogenic SIVmac239 (*8-10*). Although this unique pattern of protection is consistent with stringent CD8^+^ T_EM_-mediated control of the nascent SIV infection, epitope mapping of the 68-1 RhCMV/SIV vector-elicited CD8^+^ T cells revealed that the SIV epitopes recognized by these cells were non-overlapping with those recognized by MHC-Ia-restricted CD8^+^ T cells elicited by SIV itself or conventional SIV vaccines. Indeed, these CD8^+^ T cells were exclusively restricted by MHC-II or MHC-E, not MHC-Ia, and included recognition of universal epitopes termed supertopes (*11, 12*). Unconventionally restricted CD8^+^ T cells were not observed in RM naturally infected with RhCMV, only in RM inoculated with recombinants derived from strain 68-1 (*11, 12*). As described in detail in the companion paper (*13*) the unusual ability of 68-1 RhCMV to elicit unconventionally restricted CD8^+^ T cells is due to inversion of a genomic fragment that occurred during *in vitro* passage of this strain prior to cloning, resulting in deletion or reduced expression of eight genes in two non-contiguous gene regions (Rh157.5/.4 and Rh158-161). Complete or differential repair of these genes results in vectors inducing MHC-Ia-restricted or a mixture of MHC-Ia- and MHC-II restricted responses, the latter without supertopes. Although SIV-specific CD8^+^ T cell responses elicited by repaired vectors were similar in magnitude and memory differentiation state to responses elicited by parental 68-1 vectors, they did not protect against SIV. These results demonstrated that viral gene products that interfere with the induction of unconventional CD8^+^ T cells, including MHC-E-restricted responses, also eliminate anti-SIV efficacy. However, it is unknown what enables RhCMV to induce these responses and whether viral gene products promoting MHC-E-restricted T cells are required for efficacy.

MHC-E (HLA-E in humans, Mamu-E in RM) is generally not involved in antigen presentation to T cells but binds the conserved peptide VMAPRTL(L/V/I)L (VL9) which is embedded in the leader sequence of polymorphic MHC-Ia molecules (*14*). Similar to classical MHC-Ia, the loading of MHC-E with VL9 requires the peptide transporter associated with antigen processing (TAP) and the chaperone tapasin (*15*). Upon egress from the endoplasmic reticulum (ER) MHC-E/VL9 serves as a ligand for inhibitory CD94/NKG2A or activating NKG2C natural killer (NK) cell receptors (*16*), thus allowing NK cells to monitor pathogen-mediated or neoplastic disturbances in the classical MHC-Ia antigen presentation pathway. Interestingly, both HCMV and RhCMV encode their own VL9 peptide in the leader sequence of UL40 and Rh67, respectively (*17-19*). UL40-encoded VL9 was shown to contribute to both evasion of NKG2A^+^ NK cells by overcoming a shortage of endogenous VL9 resulting from viral MHC-Ia interference (*17, 18, 20*) and expansion of NKG2C^+^ NK cells observed in HCMV-infected individuals (*21*).

Although co-expression of Rh67 supported the intracellular transport and surface expression of an HLA-E/β2-microglobulin (β2m) single chain construct similar to UL40 (*12*), Rh67 shows very little homology to UL40 except for VL9 and adjacent amino acids (**fig. S1A**). Moreover, the Rh67 amino-terminus preceding VL9 is unrelated to UL40 and significantly shorter. For UL40 it has been demonstrated that the assembly of VL9 with HLA-E occurs even when TAP is inhibited by the HCMV US6 gene product (*17, 18*). Since TAP-inhibition is conserved in RhCMV (*22*), we examined whether Rh67 provides TAP-independent VL9 loading of Mamu-E by monitoring the intracellular transport and surface expression of Mamu-E in telomerized rhesus fibroblasts (TRF) co-transfected with the TAP-inhibitor UL49.5 of pseudorabies virus (PRV) (*23*) and a V5-epitope tagged Rh67 or UL40 (**Fig. 1A**). While Mamu-E was retained in the ER in UL49.5-expressing TRF, as indicated by sensitivity to Endoglycosidase H (EndoH), an EndoH-resistant Mamu-E population (ER “released”) was observed in the presence of either Rh67 or UL40 (**Fig. 1B**), resulting in increased surface expression of Mamu-E (**Fig. 1C, fig. S1B**). In addition to the cell surface, Mamu-E located to intracellular vesicles in Rh67- and UL40-expressing, TAP-inhibited cells, as indicated by punctate staining patterns in immunofluorescence assays (IFA), whereas Mamu-E co-localized with the ER-marker Calnexin in the absence of Rh67 or UL40 (**Fig. 1D, fig. S1C**). Transport to intracellular vesicles seemed to occur via endocytosis since MHC-E/antibody complexes were internalized from the cell surface and partially co-stained with the early endosomal marker EEA1 (**Fig. 1E, fig. S1D**). Thus, we conclude that both Rh67 and UL40 target MHC-E to the plasma membrane and then to endosomes in a TAP-independent manner.

**Figure 1.**
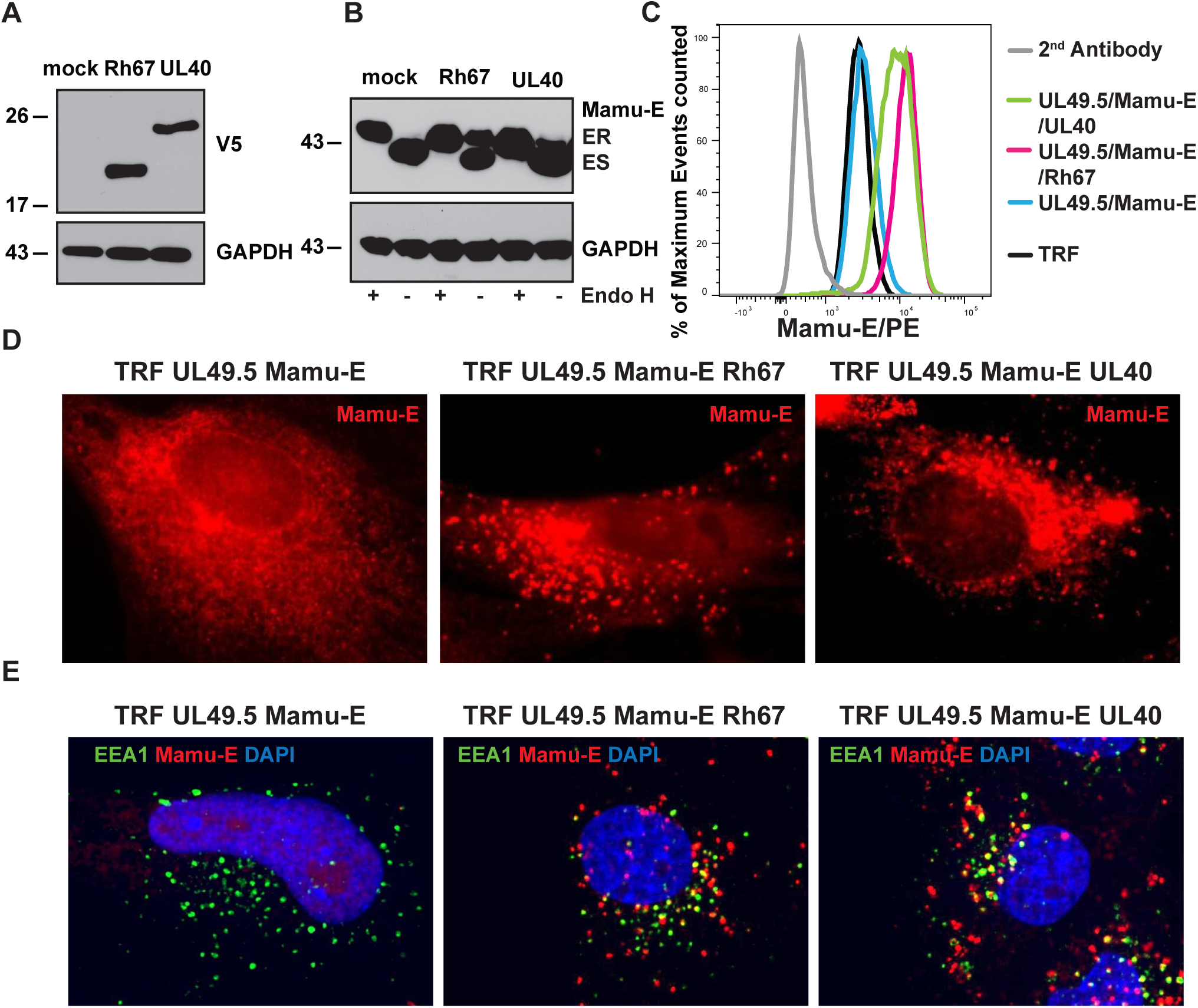
Rh67 and UL40 mediate TAP-independent intracellular maturation and internalization of MHC-E. **(A)** Expression of V5-tagged Rh67 and UL40 was determined by immunoblot in TRF that were also transduced with the TAP-inhibitor UL49.5 and Mamu-E*01:03. Anti-GAPDH was used as loading control. **(B)** Intracellular transport of Mamu-E was monitored by treating lysates of the same cell lines with Endoglycosidase H (EndoH) prior to electrophoretic separation and immunoblot with MHC-E specific antibody (ES, Endo H-sensitive; ER, Endo H-resistant). **(C)** Surface expression of Mamu-E was monitored by flow cytometry on the indicated cell lines. Rh67 and UL40 transduced cells were gated for GFP, which is co-expressed with UL49.5, and V5-staining prior to Mamu-E analysis (**fig. S1B**). **(D)** Intracellular localization of V5-taggegd Rh67 and UL40 was determined in UL49.5-expressing, Mamu-E-transfected TRF by immunofluorescence assay using indicated antibodies. Co-staining with the ER-marker Calnexin is shown in **fig. S1C. (E)** Internalization of MHC-E was monitored by adding anti-MHC-E antibody for 1 hour at 37^°^C prior to fixation, permeabilization and co-staining for Early Endosomal Antigen 1 (EEA1). Representative results are shown. Quantification of results is shown in **fig. S1D**.

To evaluate the role of Rh67-embedded VL9 in supporting Mamu-E expression in RhCMV-infected TRF, we generated a series of 68-1 RhCMV recombinants in which Rh67 was a) deleted (ΔRh67), or b) replaced with mutated Rh67, with mutations designed to truncate VL9 (deletion of the first 8 AA thus removing key N-terminal residues in the VL9 sequence; Rh67Δ1-8) or to prevent optimal VL9-MHC-E binding via replacement of VL9-position 2 anchor methionine with threonine [Rh67/M>T, a naturally occurring VL9 variant with lower affinity for MHC-E binding (*16*)], or c) replaced with its HCMV ortholog UL40 (ΔRh67/UL40) (**figs. S2A**,**B**). Since these recombinants were derived from 68-1 RhCMV/SIVgag (*5*) they all contain SIVgag as heterologous antigen (**figs. S2A**,**B**). Immunoblot analysis of Mamu-E in TRF infected with these constructs revealed that, whereas the parental 68-1 RhCMV converts the majority of Mamu-E from ER-retained to ER-released, this conversion did not occur in TRF infected with ΔRh67 or Rh67Δ1-8 (**Fig. 2A**). However, replacement of Rh67 with UL40 (ΔRh67/UL40) resulted in egress of Mamu-E indicating that UL40 can functionally replace Rh67, and replacement of Rh67 with the Rh67/M>T mutant similarly resulted in the majority of Mamu-E attaining EndoH resistance (**Fig. 2A**). The latter is in contrast to the lack of HLA-E/β2m single chain egress in cells co-transfected with Rh67/M>T (*12*), suggesting that in the context of viral infection the M>T variant supports MHC-E egress. In contrast to Mamu-E, all Rh67-derivatives were EndoH-sensitive, consistent with ER-localization, including Rh67Δ1-8 suggesting that the N-terminal truncation did not prevent protein translocation (**fig. S2C**). The requirement for viral VL9 to mediate MHC-E egress in virally infected cells is likely due to elimination of endogenous VL9 sources by viral inhibitors of MHC-Ia antigen processing encoded by Rh182, Rh185, Rh186, Rh189 and Rh178 (*19, 22, 24, 25*). To examine whether the intracellular transport of MHC-E could be restored in the absence of Rh67 in cells infected with RhCMV that also lacked all viral inhibitors of MHC-Ia presentation, we generated ΔRh67ΔRh178ΔRh182-9 (**figs. S2A**,**B**). As shown in **Fig. 2B**, we observed EndoH-resistant Mamu-E in cells infected with this vector consistent with endogenous VL9 being able to provide the MHC-E ligand in the absence of MHC-Ia inhibitors. Taken together, these data support the interpretation that Rh67 and UL40 evolved as a viral strategy to upregulate MHC-E in infected cells while at the same time preventing MHC-Ia peptide loading and surface expression.

**Figure 2.**
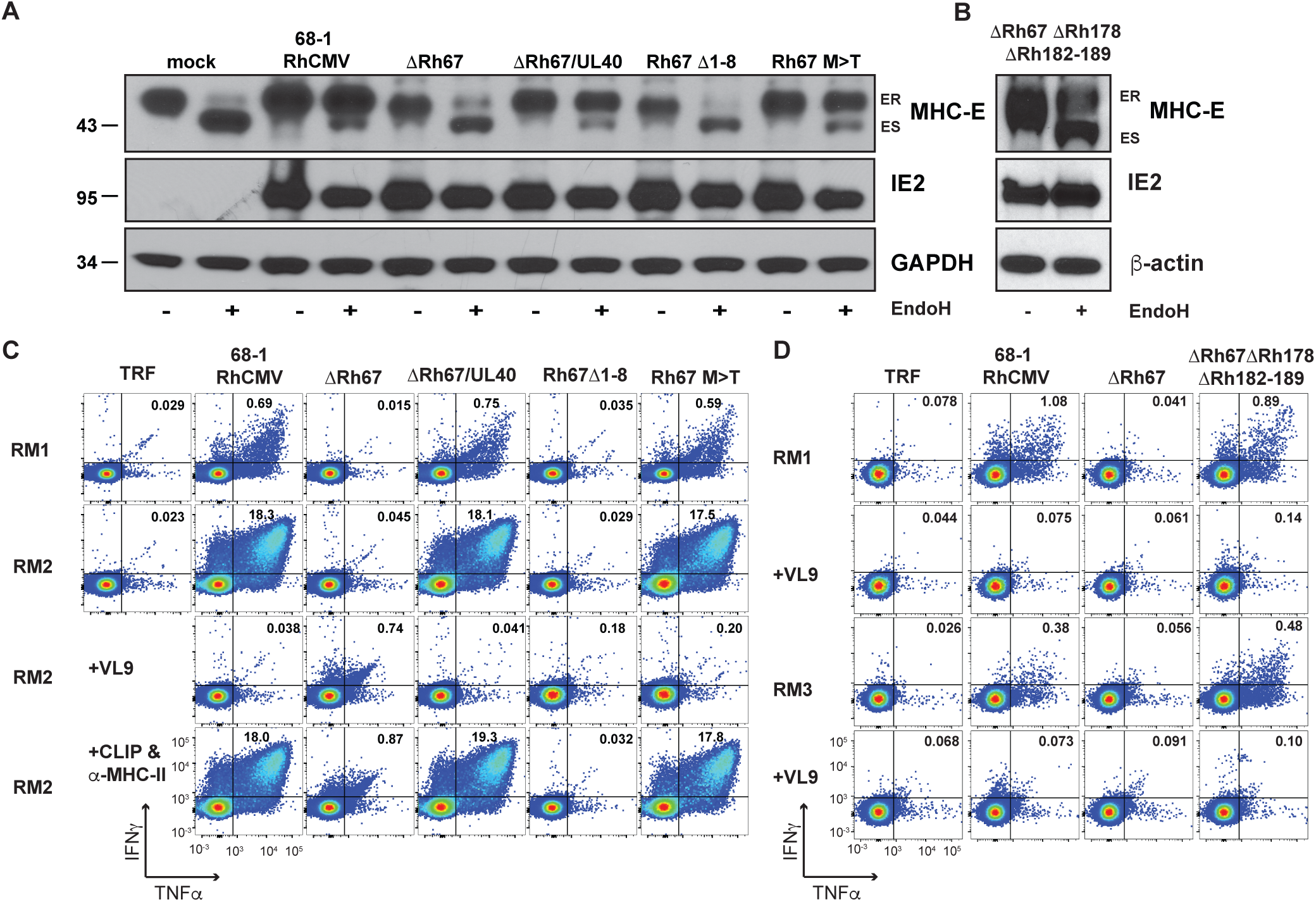
VL9-dependent intracellular transport of MHC-E is required for T cell recognition of RhCMV-infected fibroblasts. **(A**) 68-1 RhCMV/SIVgag or derivative recombinants engineered to lack Rh67 or express Rh67 lacking the first 8 amino acids of the protein (Rh67*Δ*1-8) or a Rh67 with a threonine replacing methionine at position 2 of the embedded VL9 peptide (Rh67 M>T), or with Rh67 replaced by the HCMV ortholog UL40 (*Δ*Rh67/UL40) (**fig. S2**) were used to infect TRFs at an MOI of 3. At 48 hours post-infection (hpi), cells were lysed and lysates were treated with EndoH prior to immunoblot for Mamu-E, RhCMV IE or GAPDH (ES, Endo H-sensitive; ER, Endo H-resistant). **(B)** In a separate immunoblot experiment, we infected TRF with the same MOI of 68-1 RhCMV/SIVgag deleted for Rh67, along with all inhibitors of MHC-I antigen presentation encoded by Rh178 and US6 family genes contained in the region spanning Rh182-189 (**fig. S2**). **(C)** Intracellular cytokine staining for IFN-γ and TNF of CD8^+^ T cells obtained from two 68-1 RhCMV/SIV-vaccinated RM after co-culture with uninfected TRF (negative control) or TRF infected with the indicated viruses. Where indicated, reagents for blocking MHC-E (VL9 peptide) or MHC-II (CLIP peptide and anti-MHC-II mAb G46-6) binding were added to the assay. **(D)** ICS of CD8^+^ T cells from two 68-1 RhCMV/SIV-vaccinated RM upon co-culture with the indicated viruses in the presence or absence of MHC-E-blocking VL9 peptide.

To determine whether Rh67 enables MHC-E-restricted, RhCMV-specific T cells to recognize RhCMV-infected cells, we used intracellular cytokine staining (ICS) to monitor the response of CD8^+^ T cells from 68-1 RhCMV vector-immunized RM to TRF infected with Rh67-intact vs. Rh67-modified 68-1 RhCMV. CD8^+^ T cells from these vaccinated RM include RhCMV-specific CD8^+^ T cell responses that are restricted by MHC-E and MHC-II, as well as MHC-Ia since these RM were naturally infected with wildtype RhCMV prior to 68-1 vector vaccination (*11*). However, since fibroblasts do not express MHC-II and since the MHC-Ia alleles expressed by the TRFs used for infection would be both allogeneic to the CD8^+^ T cells and effectively downregulated by infection (*19, 22, 24*), we expected that most recognition would be MHC-E-restricted. Indeed, essentially all CD8^+^ T cell recognition of 68-1 RhCMV-infected TRF was blocked by exogenous VL9 peptide, but not by blocking of MHC-II, consistent with dominant MHC-E-restricted epitope presentation (**Fig. 2C**). In contrast, the same CD8^+^ T cells did not recognize TRFs infected with *Δ*Rh67 or Rh67Δ1-8, but did recognize TRFs infected with 68-1 RhCMV when Rh67 was replaced by its HCMV ortholog UL40 or with Rh67/M>T (**Fig. 2C**). Interestingly, Rh67 was not needed for MHC-E-restricted recognition if TRF were infected with ΔRh67ΔRh182-9ΔRh178 (**Fig. 2D**). Since this vector lacked all viral inhibitors of MHC-Ia presentation, infected TRF were also recognized by MHC-Ia-matched SIV Gag epitope-specific CD8^+^ T cells, in contrast to 68-1 RhCMV or ΔRh67 which both prevent antigen presentation by MHC-Ia (**fig. S3**). We conclude that VL9-induced intracellular transport of Mamu-E is essential for the presentation of non-VL9 peptides to MHC-E-restricted CD8^+^ T cells, with viral VL9 needed when sufficient endogenous VL9 is not available.

We next asked whether *in vivo* priming of MHC-E-restricted CD8^+^ T cells had similar requirements for VL9-induced Mamu-E trafficking by comparing the SIV Gag-specific CD8^+^ T cell responses in RM immunized with Rh67-intact vs. -deleted 68-1 RhCMV/SIVgag. Importantly, the ΔRh67 vector elicited robust overall SIV Gag-specific T cell responses comparable to those of the parent Rh67-intact 68-1 vector (**Fig. 3A**), indicating that, unlike viral inhibitors of MHC-Ia antigen presentation (*24*) or viral inhibitors of NKG2D-dependent NK cell activation (*26*), Rh67 was not required to establish immunogenic infection, even in RMs with pre-existing anti-RhCMV immunity. However, lack of Rh67 had a profound influence on the epitopes targeted by the SIV Gag-specific CD8^+^ T cells in the ΔRh67-vaccinated RM. Although these CD8^+^ T cells recognized MHC-II-restricted supertopes, they did not respond to the MHC-E-restricted supertopes (**Fig. 3A**). Full epitope deconvolution followed by restriction analysis revealed that the entire response was MHC-II-restricted suggesting that MHC-E-restricted T cell priming was abrogated. Expression of UL40 instead of Rh67 (ΔRh67/UL40) restored MHC-E-restricted CD8^+^ T cell priming, but neither the Rh67Δ1-8 nor Rh67/M>T vectors were able to prime these responses (**Figs. 3B**). Moreover, in the absence of Rh67, restoration of endogenous VL9 production by deletion of viral inhibitors of MHC-Ia presentation (ΔRh67ΔRh178ΔRh182-9) was unable to support MHC-E-restricted CD8^+^ T cell priming. Due to deletion of Rh189 (homolog of US11 in HCMV), the ΔRh67ΔRh178ΔRh182-9 vector elicited CD8^+^ T cell responses to canonical MHC-Ia epitopes, (*11*), which resulted in mixed recognition of MHC-II-and MHC-Ia-restricted epitopes (**fig. S4**). Thus, regardless of viral interference with antigen presentation by MHC-Ia, induction of MHC-E-restricted CD8^+^ T cell responses requires viral provision of VL9 by a Rh67/UL40-mediated process. This result also indicates that these responses must arise from direct presentation by Rh67/UL40-expressing, 68-1 RhCMV vector-infected cells, and that direct priming depends more stringently on viral VL9 peptide than infected cell recognition *in vitro*.

**Figure 3.**
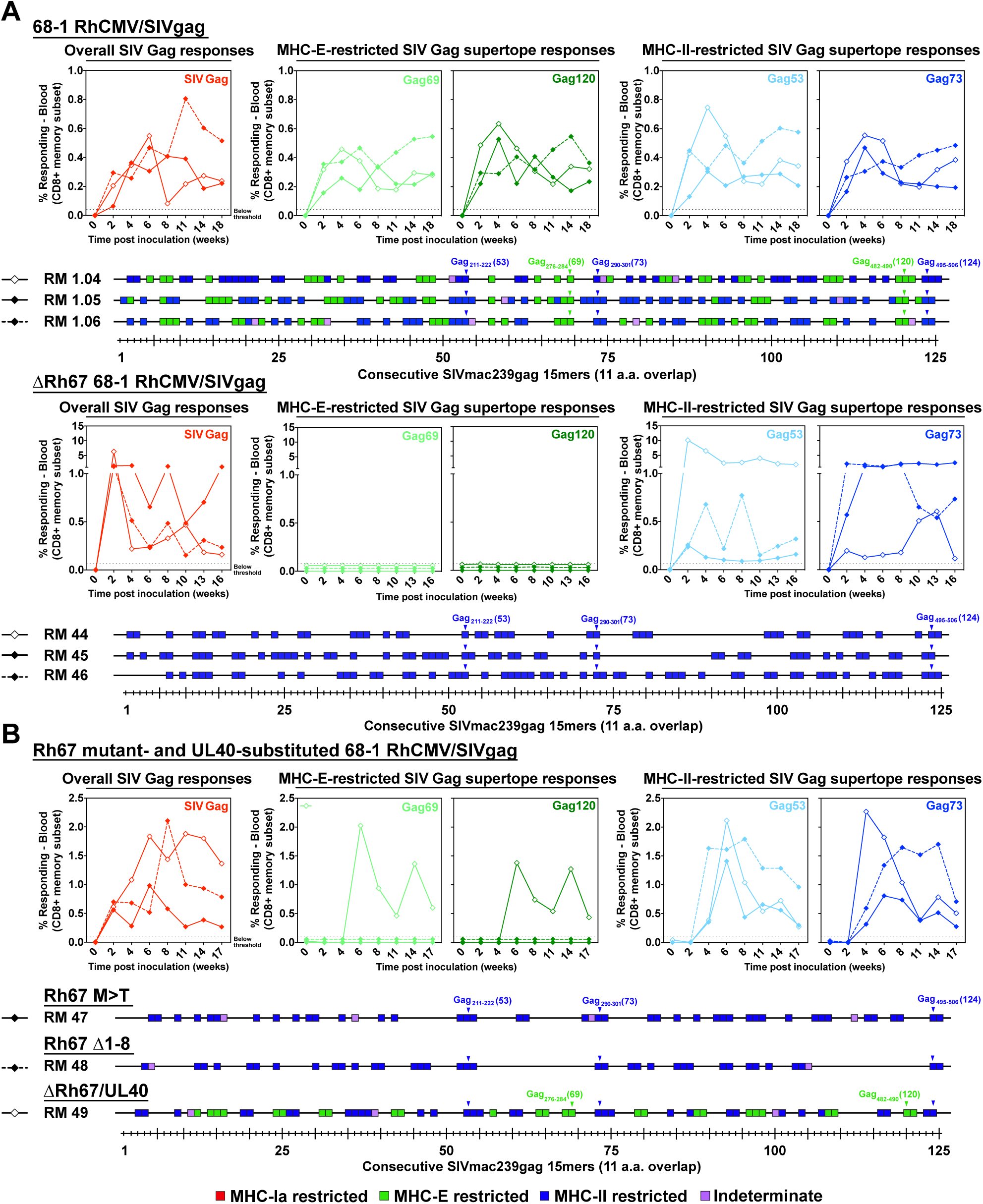
Role of Rh67 in RhCMV/SIV vector CD8^+^ T cell response programming. **(A)** Comparison of SIV Gag-specific CD8^+^ T cell responses elicited by 68-1 RhCMV/SIVgag (RMs 1.04, 1.05, 1.06) vs. ΔRh67 68-1 RhCMV/SIVgag (RMs 44, 45, 46) vectors. Peripheral blood CD8^+^ T cells from these RM inoculated with each of these vectors were assessed by a flow cytometric intracellular cytokine assay for responses to a mixture of 125 consecutive 15mer peptides comprising the SIV Gag protein sequence (red; top panel), to individual MHC-E and MHC-II-restricted SIV Gag supertopes (green and blue, respectively; top panel), and to each of 125 consecutive 15mer SIV Gag peptides with any above threshold (≥ 0.05% after background subtraction) responses indicated by a box. Boxes are colored to reflect MHC restriction based on the ability to inhibit the response with the MHC-E blocking peptide VL9, the MHC-II blocking mAb G46-6, or the pan-MHC-I blocking mAb W6/32 (*13*). In the bottom panel, arrowheads indicate the15mer peptides including the supertopes designated in the top panel. (**B**) Similar analysis of SIV Gag-specific CD8^+^ T cell responses elicited by Rh67 M>T (RM47), Rh67*Δ*1-8 (RM 48), or *Δ*Rh67/UL40 (RM49) 68-1 RhCMV/SIVgag vectors.

To determine whether MHC-E-restricted CD8^+^ T cell priming is required for 68-1 RhCMV/SIV vaccine efficacy, we vaccinated 15 RM with three ΔRh67 68-1 RhCMV/SIV vectors individually expressing SIVgag, SIVretanef (a fusion of rev, tat and nef sequences), or a 5’ fragment of SIVpol [comparing the immunogenicity and efficacy to assessment of the Rh67-intact parent 68-1 RhCMV/SIV vectors in the companion manuscript (*13*) (**fig. S5**)]. Importantly, Rh67 deletion did not diminish the magnitude of the overall vector-elicited, SIV-specific T cell responses (indeed, SIV-specific CD4^+^ T cell responses were significantly more robust in RM vaccinated with Rh67-deleted vector) or substantially alter either the highly T_EM_-biased response phenotype or the cytokine response pattern compared to parental 68-1 (**Fig. 4A-E**). However, as expected, the *Δ*Rh67 68-1 RhCMV/SIV vector-elicited CD8^+^ T cell response to all SIV proteins was entirely MHC-II-restricted, including MHC-II-restricted supertopes (**Figs. 4C; fig. S6**). Next, we performed repeated, limiting dose, intrarectal SIVmac239 challenge on this cohort, as previously described (*8-10, 13*). Whereas 8 of 15 RM vaccinated with parental 68-1 RhCMV/SIV vectors showed typical RhCMV/SIV vaccine efficacy (induction of SIVvif-specific T cell responses and development of cell-associated viral loads in tissues, in the absence of sustained plasma viremia), 14 of the 15 ΔRh67 68-1 RhCMV/SIV vector-vaccinated RM showed progressive SIV infection identical to unvaccinated controls (**Fig. 4F**). One of 15 vaccinated RM showed restricted viremia (peaking at 6,900 SIV RNA copies/ml) before controlling viremia to below the detection limit by week 4 post infection. Since the pattern of this viral control is typical of conventional elite control (and atypical for RhCMV/SIV vaccine-mediated replication arrest), we treated this RM with a CD8*α*-targeted depleting monoclonal antibody (mAb) at 8 weeks post-infection (**Fig. 4G**). Previous observations established that 68-1 RhCMV/SIV vector-mediated replication arrest is not abrogated by CD8*α*^+^ cell depletion (*8, 9*), even when cell-associated virus is readily demonstrable in tissues (**fig. S7**), whereas conventional elite control is universally, if transiently, abrogated by this treatment (*27*). As shown in **Fig. 4F**, this RM’s viral control was temporarily abrogated by the anti-CD8*α* treatment, confirming that this RM is a conventional elite SIV controller (with control therefore unrelated to RhCMV/SIV vaccination-mediated replication arrest). Thus, we conclude that ΔRh67 68-1 RhCMV/SIV vaccine lacked efficacy against SIV in this challenge study, and that the MHC-II-restricted CD8^+^ T cells elicited by this vaccine are not, at least by themselves, able to mediate SIV replication arrest.

**Figure 4.**
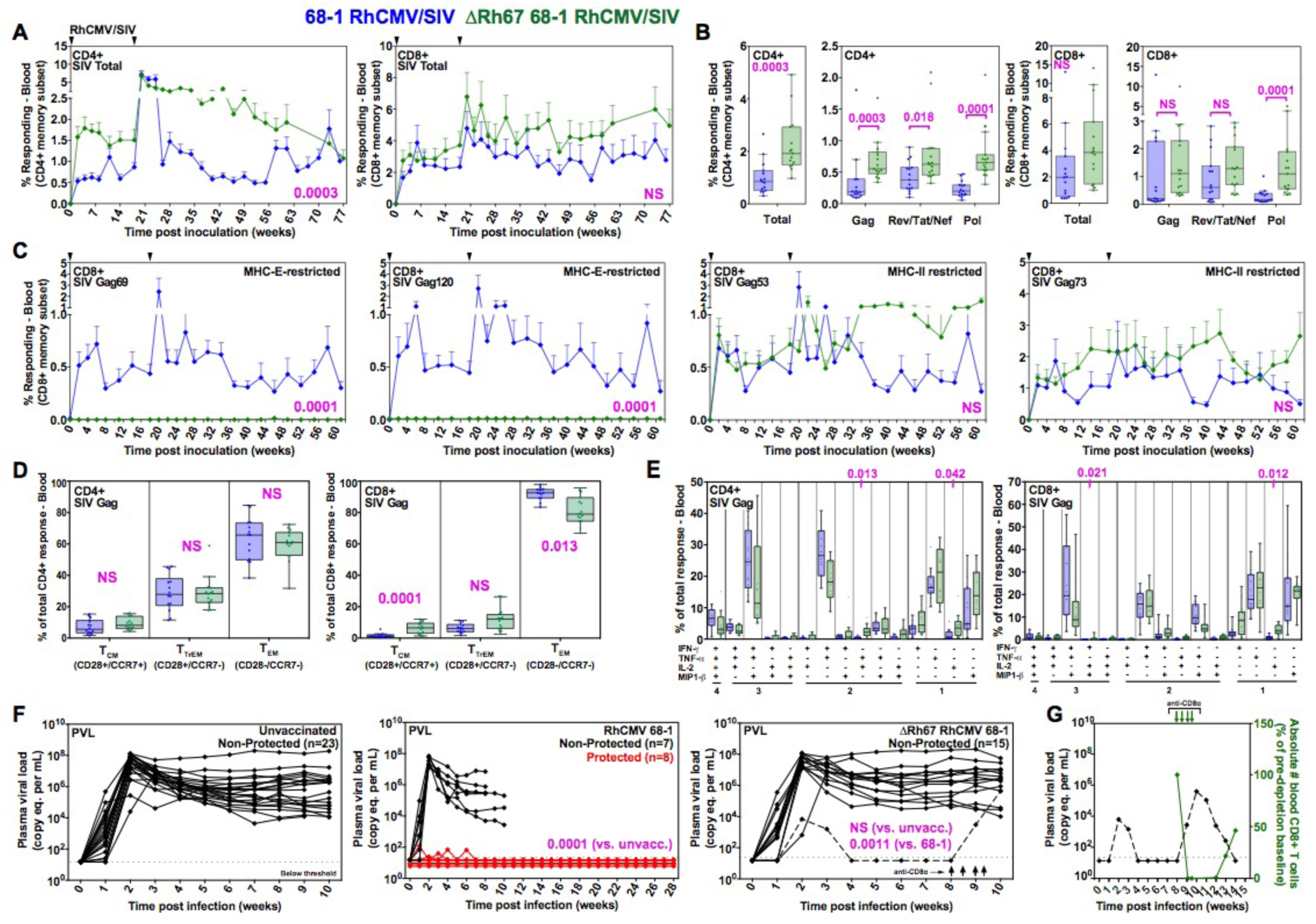
Immunogenicity and efficacy of ΔRh67 68-1 RhCMV/SIV vectors. **(A**,**B)** Longitudinal and plateau-phase analysis of the vaccine-elicited SIV Gag-, Rev/Tat/Nef-, and Pol-specific CD4^+^ and CD8^+^ T cell responses in peripheral blood of the RM vaccinated with a ΔRh67 68-1 RhCMV/SIV vector set compared to the Rh67-intact parent 68-1 vector set (see **Fig. S5**). In **A**, the background-subtracted frequencies of cells producing TNF-*α* and/or IFN-*γ* by flow cytometric ICS assay to peptide mixes comprising each of the SIV inserts within the memory CD4^+^ or CD8^+^ T cell subsets were summed for overall responses with the figure showing the mean (+ SEM) of these overall responses at each time point. In **B**, boxplots compare the total and individual SIV insert-specific CD4^+^ and CD8 ^+^ T cell response frequencies between the two vaccine groups during the vaccine phase plateau. (**C**) Longitudinal analysis of the vaccine elicited CD8^+^ T cell responses to SIV Gag supertopes in peripheral blood of each vaccine group by ICS assay. Gag_276-284_ (69) and Gag_482-490_ (120) are MHC-E-restricted supertopes (*12*); Gag_211-222_ (53) and Gag_290-301_ (73) are MHC-II-restricted supertopes (*11*). (**D**) Boxplots compare the memory differentiation of the vaccine-elicited CD4^+^ and CD8^+^ memory T cells in peripheral blood responding to overall SIV Gag 15mer peptide mix with TNF and/or IFN-*γ* production during the vaccine phase plateau. Memory differentiation state was based on CD28 and CCR7 expression, delineating central memory (T_CM_), transitional effector-memory (T_TrEM_), and effector-memory (T_EM_), as designated. (**E**) Boxplots compare the frequency of vaccine-elicited CD4^+^ and CD8^+^ memory T cells in peripheral blood during plateau phase responding to the overall SIV Gag 15mer peptide mix with TNF, IFN-*γ*, IL-2, and MIP-1*β* production, alone and in all combinations. Wilcoxon p-values for comparison of all T cell response parameters shown in panels A-E are shown where significant (p<0.05; adjusted for multiple comparisons in panels B-E). (**F**,**G**) Assessment of the outcome of SIV infection after repeated, limiting dose SIV_mac239_ challenge of the ΔRh67 68-1 RhCMV/SIV vector vaccinated RM, relative to RM vaccinated with the Rh67-intact parent 68-1 vectors [red delineates RM with documented SIV replication arrest; reproduced from (*13*)], and to unvaccinated controls (including RM contemporaneously challenged with both vaccinated cohorts). One of 15 ΔRh67 68-1 RhCMV/SIV vector set vaccinated RM showed abbreviated SIV viremia post-infection prompting us to perform CD8*α* cell depletion at day 57 post-infection to distinguish conventional elite control (susceptible to CD8*α* cell depletion) to RhCMV/SIV vector vaccination induced SIV replication arrest (not susceptible to CD8*α* cell depletion; **fig. S7**). Binomial exact p-values are shown where the proportion of RM with SIV replication arrest (protected; shown in red) in one vaccine group differs significantly from the designated comparison group. NS = not significant.

In the companion manuscript (*13*), we demonstrate that RhCMV/SIV vectors eliciting MHC-Ia-restricted CD8^+^ T cell responses or a mixture of MHC-Ia- and MHC-II-restricted CD8^+^ T cell responses (the latter without MHC-II supertope responses) are also non-efficacious. Taken together, these data strongly suggest that MHC-E-restricted CD8^+^ T cell responses are required for protection. Since all RhCMV/SIV vector-elicited CD8^+^ T cell responses appear to be phenotypically and functionally similar (**Fig. 4**) (*13*), the requirement for MHC-E-restricted CD8^+^ T cell responses for anti-SIV efficacy is most likely attributable to advantages afforded by MHC-E-mediated CD8^+^ T cell recognition of SIV-infected cells in early SIV infection, possibly due to the upregulation of MHC-E in SIV-infected CD4^+^ T cells relative to SIVnef-mediated down-regulation of MHC-Ia (*12, 28*) and limited MHC-II expression on the resting CD4^+^ T cells that are dominant early viral targets (*29*). SIV appears to have evolved to evade NK and MHC-Ia-restricted CD8^+^ T cells in the absence of selection pressure from MHC-E-restricted CD8^+^ T cells since these cells are not naturally primed during SIV infection (*12*). This is in contrast to the more complex immune evasion strategy of RhCMV, which efficiently prevents antigen presentation by MHC-Ia via the Rh178 and Rh182-189-encoded proteins (*19, 22, 24, 25*) (**fig. S3**), while simultaneously enhancing the intracellular transport of MHC-E via the VL9-providing Rh67 protein (**Fig. 2**), a process that in the absence of specific inhibition leads to MHC-E-restricted CD8^+^ T cell priming. Indeed, the presence of eight different genes that each prevent MHC-E-restricted CD8^+^ T cell priming in the RhCMV genome [Rh157.5/.4 and Rh158-161] (*13*), strongly suggests that the efficient NK cell evasion afforded by the Rh67 mechanism of MHC-Ia-independent MHC-E upregulation comes at a price of increased recognition by MHC-E-restricted CD8^+^ T cells, which, in turn, is countered by these viral-encoded priming inhibitors.

It should, however, be noted that the finding that Rh67 function leads to MHC-E-restricted T cell priming is counter intuitive since VL9 binds with higher affinity to MHC-E than non-VL9 peptides and thus might be expected to prevent or displace binding of unrelated, lower affinity peptides (*30, 31*). Instead, our data suggest that while loading with viral VL9 is required for MHC-E egress from the ER in the absence of host VL9, there is in place a host mechanism that provides for exchange of the VL9 peptide with other viral peptides resulting in MHC-E-mediated antigen presentation to CD8^+^ T cells. This model is supported by the correlation of MHC-E egress with T cell recognition of RhCMV vector-infected cells and by the finding that surface MHC-E traffics to intracellular vesicles. The exact nature of these vesicles still needs to be determined, but our observation is consistent with previous reports showing that, in monocytes, HLA-E co-localized with markers of endolysosomal and autophagosomal vesicular structures (*32*). Moreover, structural and biochemical studies demonstrate that VL9 bound to HLA-E can be exchanged for lower-affinity, non-VL9 peptides most likely due to an unusual stability of “empty” MHC-E/β2m heterodimers (*12, 30*). Thus, with this novel epitope-loading mechanism, the host partially counters the viral NK cell evasion strategy by eliciting a novel MHC-E-restricted CD8^+^ T cell response which would preferentially recognize CMV-infected cells with Rh67-mediated MHC-E up-regulation. Taken together, these data suggest that 68-1 RhCMV/SIV vector efficacy results from a mismatch in immune evasion strategies between RhCMV and SIV, with the former virus able to generate novel responses that it has evolved to evade, but the latter virus cannot. Although HCMV and RhCMV are different viruses, they do share a close evolutionary relationship and the relevant mechanisms appears to be conserved (*28*), including the ability of UL40, UL128/UL130 and UL146/147 to functionally replace their RhCMV orthologs (*13*), offering the hope that a 68-1-like HCMV-based vaccine vector might be able to take advantage of the same mismatch to provide an effective HIV/AIDS vaccine.

## ACKNOWLEDGMENTS

We thank T. Whitmer, A. Bhusari A. Legasse, M. Fischer, C. Shriver-Munsch, T. Swanson, A. Sylwester, S. Hagen, E. McDonald, K. Randall, and K. Rothstein for technical or administrative assistance; B. Keele (Frederick National Laboratory) for providing SIVmac239 challenge virus, and A. Townsend for figure preparation. We acknowledge the sequencing services of the OHSU Massively Parallel Sequencing Shared Resource and the ONPRC Molecular Technologies Core. We also thank the ONPRC Molecular Virology Core for virus stock preparation.

## AUTHOR CONTRIBUTIONS

KF and LJP conceived the experimental strategy, supervised experiments, analyzed and interpreted data, and wrote the paper. SGH planned and performed animal experiments and immunologic assays, assisted by RMG, CMH, ABV, JCF and AS. MV, RI and NJ generated and characterized transfected cell lines. D. Malouli, JW and IS constructed and quality-tested recombinant RhCMV constructs. MV, RI and D. Morrow performed in vitro T cell assays with recombinant RhCMV constructs. KO, RS, BB, WJB, MH and JDL performed SIV plasma viral load quantifications. MKA managed the animal care and procedures. PTE planned and performed all statistical analyses, assisted by JS.

## DATA AND MATERIALS AVAILABILITY

All data associated with this study are present in the paper or Supplementary Materials. RhCMV/SIV vectors can be obtained through an MTA.

