## Supplemental Material for "Modulation of MHC-E transport by viral decoy ligands is required for RhCMV/SIV vaccine efficacy"

**SUPPLEMENTARY MATERIALS LIST:**

**Materials and Methods**

**Figure S1.** Rh67 and UL40 contain a conserved VL9 sequence and cause TAP-independent Mamu-E transport and internalization

**Figure S2.** Description and characterization of Rh67-modified 68-1 RhCMV/gag recombinants.

**Figure S3.** Triggering of Mamu-A01-restricted, SIVgagCM9-specific, CD8$^+$ T cells by RhCMV recombinants.

**Figure S4.** Requirement for Rh67 for MHC-E-restricted CD8$^+$ T cell priming in the absence of all viral inhibitors of MHC-Ia presentation.

**Figure S5.** Protocols for the comparison of the immunogenicity and efficacy of Rh67-deleted vs. Rh67-intact RhCMV/SIV vectors.

**Figure S6.** Epitope analysis of ΔRh67 68-1 RhCMV/SIV vaccine-elicited, SIV-specific CD8$^+$ T cell responses in RMs destined for SIV challenge.

**Figure S7.** Resistance of 68-1 RhCMV/SIV vaccine-induced replication arrest to *in vivo* CD8α$^+$ cell depletion.

### MATERIALS AND METHODS

**Study Design.** The objectives of this study were to a) evaluate the role of Rh67-encoded MHC-E ligands on the ability of 68-1 RhCMV vectors to present MHC-E-restricted epitopes *in vitro*, b) elicit MHC-E-restricted CD8$^+$ T cell responses *in vivo,* and c) determine whether vaccination of RM with RhCMV/SIV vectors lacking Rh67 would provide the same stringent post-acquisition control of SIVmac239 infection as Rh67-intact 68-1 RhCMV/SIV vectors. *In vitro* studies were performed with life-extended primary rhesus fibroblasts infected with RhCMV to determine the impact of Rh67-deletions or –modifications on the ability to stimulate MHC-E-restricted CD8$^+$ T cells. To minimize the number of RM used in these experiments, most were designed with the goal to generate qualitative rather than quantitative comparisons with Rh67-intact vectors. Because we observed highly consistent results and simultaneously monitored multiple epitopes as few as one animal per construct was sufficient to determine whether a given Rh67-modified RhCMV vector was able to elicit MHC-E restricted CD8$^+$ T cells. To evaluate ΔRh67 68-1 RhCMV/SIV vector immunogenicity and efficacy we appended analysis of this vector backbone to the large NHP study described in the companion manuscript comparing immunogenicity and efficacy of Rh157.5/157.5 gene region-modified 68-1 RhCMV/SIV vectors with the parental 68-1 RhCMV/SIV vector (*13*). We constructed and validated a 3-vector set expressing the same SIV Gag, Rev/Nef/Tat and 5'-Pol inserts as in the companion study and subcutaneously administrated these vectors to 15 randomly selected male RM from the same pool as the prior study, using the identical vaccination regimen. In addition, these RM were SIV challenged with the same, repeated, limiting dose, intra-rectal challenge protocol using the same SIVmac239 stock as the companion study, side-by side with an additional 8 unvaccinated control RM (which validated the effectiveness of the challenge protocol). Since the immunogenicity and efficacy of the parental 68-1 RhCMV/SIV vectors is highly consistent across multiple studies (*8-10, 13*), we used prior analysis of the parental 68-1 RhCMV/SIV vector in the companion study (*13*) as the positive control group for this study. As previously noted (*13*), the n=15 group size allows us to detect per-vaccine-group protection levels of 14% at 90% power without multiplicity adjustment. All results from these experiments are included in the presented associated data (no data were excluded as outliers). Plasma viral load assays were assayed by blinded analysis; however, due to logistical constraints, other staff were not blinded to treatment assignments. Primary data are reported in data files 1-3.

**Cell lines.** Telomerized rhesus fibroblasts (TRF) were transduced with retrovirus pLZRS IRES GFP expressing PRV UL49.5 in front of an internal ribosomal entry site that is followed by enhanced green fluorescent protein (GFP) (*33*). After sorting for GFP expression, cells were transduced with Mamu-E*02:04 expressing lentivector pLVX EF1α IRES blasticidin (derived from pLVX EF1α IRES puro (Clontech) by replacing the puromycin resistance with blasticidin resistance cassette). After selection for blasticidin resistance the cells were transduced with pLVX EF1α IRES puro (Clontech) carrying synthetic genes encoding C-terminally V5-epitope tagged RhCMV Rh67 or HCMV UL40 (Life Technologies). Upon selection for puromycin resistance the resulting cell lines were maintained in DMEM supplemented with 10% fetal bovine serum (FBS), 100 U/ml penicillin, and 100 μg/ml streptomycin (PenStrep) with added puromycin (Invivogen, 4μg/ml) and blasticidin (Invivogen, 20μg/ml). Expression of GFP, Rh67, UL40, and Mamu-E was confirmed by immunoblot and flow cytometry **(Fig. 1, fig. S1).**

**Generation and testing of recombinant RhCMV vectors.** All recombinant viruses used in this study were generated using bacterial artificial chromosome (BAC) mutagenesis of BAC-cloned RhCMV strain 68-1 (*34*). 68-1 RhCMV expressing SIVgag was generated as previously described (*5*). This vector was used to generate Rh67-modified recombinants by homologous recombination of linear DNA fragments generated by gene synthesis (Life Technologies) with all constructs containing an in-frame fusion of the parainfluenza V5 epitope tag (GKPIPNPLLGLDST) at their C-terminus: Rh67, UL40, Rh67M>T,

Rh67Δ1-8 (**fig. S2A**). DNA fragments were cloned into pORI and amplified by PCR together with a Kanamycin-resistance (KanR) marker flanked by flippase recognition target (FRT) sites (*35*). PCR primers included 50 bp homologous to the Rh67 flanking region so that bacteriophage Lambda recombination enzymes Red α,β,γ-dependent homologous recombination resulted in exchange of Rh67 with the Rh67-modified sequence. The KanR resistance cassette that was subsequently removed by induction of the yeast flippase resulting in a remaining FRT "scar" (**fig. S2A**). Rh67 was deleted from 68-1 RhCMV/SIVgag by replacing Rh67 with the excisable KanR cassette using the same homologous recombination strategy. Rh67-deleted RhCMV/rtn and RhCMV/pol5' were generated by using homologous recombination to replace Rh67 with genes encoding the SIV antigens Rev/Tat/Nef or the 5'-fragment of SIVpol together with the excisable KanR gene.

To generate ΔRh67ΔRh178ΔRh182-9 we transformed the 68-1 RhCMV/SIVgag BAC into *E.coli* strain GS1783 to allow *en passant* recombination (*36*). Using this scarless recombination technique we sequentially deleted Rh67, Rh178, and the entire Rh182-Rh189 gene region. Each of the intermediate steps were controlled by XmaI restriction digest and Sanger sequencing across the deletions.

The final BACs were analyzed by next generation sequencing of the entire genome on an Illumina MiSeq sequencer. Consensus sequences were deposited in GenBank: ΔRh67/gag (MN622881), ΔRh67/pol5' (MN622882), ΔRh67/rtn (MN622883), ΔRh67/UL40/gag (MN622884), Rh67M>T/gag (MN622885), Rh67V5/gag (MN622886), Rh67Δ1-8/gag (MN622887), ΔRh67ΔRh178ΔRh182-9/gag (MT068444).

Recombinant viruses were reconstituted by electroporation of BAC DNA into primary rhesus fibroblasts. Briefly, cells were trypsinized, pelleted, and resuspended in Opti-MEM (Invitrogen). BAC DNA was added, and cells were pulsed at 0.25 kV, 0.95 mF using the Gene Pulser II (Bio-Rad). Upon recovery of recombinant RhCMV, the vectors were propagated in TRF. For viral stocks, virus was purified by ultracentrifugation through a 20% sorbitol cushion. The loxP-flanked BAC cassette encodes a Cre recombinase under control of a mammalian promoter resulting in spontaneous excision of the BAC cassette (*34*). Expression of SIV antigens and Rh67 constructs was confirmed by immunoblot following infection of TRF (**fig. S2B**).

**Polyacrylamide gel electrophoresis and immunoblot analysis**. Cells were harvested with trypsin/EDTA (Corning 25-051-CI) and washed with PBS. The cell pellets were then resuspended at $1 \times 10^7$ cells/ml in water with 1% NP-40 and HALT protease inhibitors (Thermo Fisher) and incubated at 4°C for 2h to lyse the cells. The resulting lysates were centrifuged at 15,000 RPM at 4°C for 45 min. The supernatants were harvested and frozen at -20°C until use. For Endo H treatment, 9µl of each lysate sample was mixed with 1.09µl of 10X glycoprotein dissociation buffer (New England Biolabs) and boiled for 10 min. The lysates were then mixed with 5.97µl of EndoH reaction buffer (50mM sodium acetate, 0.5mg/ml BSA, 1.25% NP-40 and HALT protease inhibitor) and 1.3µl of EndoH (Roche), or water for mock-treated samples, and incubated at 37° overnight. To detect a reduction in the apparent molecular weight due to removal of asparagine-linked glycans by EndoH we separated lysates on 10% polyacrylamide gels and transferred to PVDF membranes (Millipore). Following blocking with 5% milk in PBS-0.1% Tween-20 (Fisher Scientific) (PBS-T), membranes were probed with the following antibodies in 5% milk in PBS-T: anti-MHC-E (1:1000, Clone 4D12, LSBio #C179742), anti-βactin (1:10,000, Clone AC-74, Sigma #A2228) anti-FLAG (1:5000, Sigma #F3165), anti-V5 (1:500, Invitrogen #37-7500), anti-GAPDH (1:5000, Invitrogen #MA5-15738), or anti-RhCMV IE2 (*37*) (Clone IIA5.2, 1:100). Membranes were washed three times with PBS-T and then incubated with goat anti-mouse HRP (1:5000, Invitrogen #A28177) secondary antibody in 5% milk in PBS-T. Membranes were incubated with Supersignal West Pico chemiluminescent substrate (Thermo Scientific) and exposed to chemiluminescent film (GE Healthcare) or scanned on a BioRad ChemiDoc MP imager.

**Immunofluorescence.** TRF expressing Mamu-E and UL49.5 alone or together with V5-tagged Rh67 or UL40 were plated on 12mm wide, 0.13-0.17mm thick glass coverslips in 24-well plates and stained at 24 hrs post-plating. Cells were rinsed with PBS and fixed with 4% paraformaldehyde (PFA) in PBS for 10 min at room temperature (RT) followed by washing 3x with PBS, blocking with 1% BSA (Fisher) in PBS and permeabilization with 0.2% saponin (Millipore) for 1 hr at RT. Fixed cells were incubated with the following antibodies for 1 hr at RT: anti-V5 (1:400, Novus biologicals #NB600-381), anti-Calnexin (1:250, LS Bio #LS-B12410), anti-MHC-E (1:100, HLA-E, MBL #K0215-3) followed by washing 3x for a total of 30 mins using saponin-BSA buffer. Secondary antibodies (Invitrogen #A-21081, #A-11032 and #A32733) were used in 1:500 dilution in saponin-BSA buffer for 1 hr at RT. Cells were washed 3x for a total of 30 mins using saponin-BSA buffer followed by a final wash in PBS. Coverslips were mounted on glass slide using Prolong Diamond (Invitrogen) and cured for 24 hrs. The cells were visualized and imaged using 100x oil objective in Keyence BZ-X700 microscope. Images were processed and overlaid using FIJI software.

Internalization of HLA-E was monitored 24 hrs after cells were plated on 12mm wide, 0.13-0.17mm thick glass coverslips in 24-well plates by adding anti-MHC-E antibody (MBL #K0215-3; 1:100 diluted in DMEM, 2.5% FBS (D2.5%)) to TRF expressing Mamu-E and UL49.5 alone or together with Rh67-V5 or UL40-V5. After incubation at 37°C for 1 hr the cells were washed 3x in cold D2.5% and fixed with 4% PFA in PBS for 10 mins at RT. The cells were washed 3x using PBS, blocked and permeabilized with 0.2% saponin and 1% BSA buffer for 1 hr at RT. Permeabilized cells were incubated with EEA1-specific antibody (1:100, CST, 2411S) for 1 hr at RT followed by 3x washing for a total of 30 mins using saponin-BSA buffer. Secondary antibodies (1:500 in saponin-BSA buffer; Invitrogen, A-11032 and A32733) together with DAPI (1µg/ml, Invitrogen, D1306) were incubated for 1 hr at RT. Cells were washed 3x for a total of 30 mins using saponin-BSA buffer followed by a final wash in PBS. Coverslips were mounted on glass slides using Prolong Diamond (Invitrogen, P36961) and cured for 24 hrs. The cells were visualized and imaged using 100x oil objective in Keyence BZ-X700 microscope. To acquire high resolution images the 2D structured illumination feature was used. Images were processed and overlaid using FIJI software. For quantification images acquired by Keyence microscope using either the 100x or 40x objective from three independent experiment sets were compiled. The FIJI macro was used to count the nuclei and fluorescent signals were counted manually.

**Flow cytometric analysis of MHC-E expression on TRF:** TRF or TRF expressing Mamu-E and UL49.5 alone or together with Rh67 or UL40 were harvested with trypsin/EDTA (Corning) and washed with cold DMEM supplemented with 2.5% FBS (D2.5%). This media was used throughout for washing and MHC-E antibody incubation of live TRF. $2.5 \times 10^5$ cells were incubated with anti-MHC-E antibody (MBL #K0215-3, 1:100) for 1 hr at 4°C. The samples were washed 3x and incubated with phycoerythrin (PE)-conjugated secondary antibody (Thermo Fisher #31862, 1:250) and Live/Dead stain (1:100, Life Technologies #L34957) for 45 mins at 4°C. The cells were washed 3x and fixed with 4% PFA in PBS for 10 min at RT, followed by 3x washing using PBS. Cells were further stained for intracellular V5-epitope expression, after blocking with 1% BSA and permeabilization with 0.2% saponin in PBS for 1 hr at RT. Anti-V5 antibody (1:400, Novus biologicals #NB600-381) was incubated in saponin-BSA buffer for 1 hr at RT and cells were washed 3x using saponin-BSA buffer. The samples were washed 3x and incubated with anti-rabbit secondary antibody conjugated with Alexa fluor 647 (Thermo Fisher #A-21244, 1:500). Finally, stained cells were fixed in 1% PFA in PBS and analyzed on a BD LSRII Cytometer.

**Rhesus macaques.** The experiments reported here used a total of 65 purpose-bred male and female RM (*M. mulatta*) of Indian genetic background, including 3 RM previously immunized with 68-1 RhCMV vectors for separate studies but used here as source of MHC-E-restricted $CD8^+$ T cells, 7 RM for immunogenicity analysis of 68-1-based, Rh67-variant RhCMV/SIVgag vectors, 53 RM for comparative immunogenicity and efficacy analysis of Rh67-intact vs. ΔRh67 68-1 RhCMV vectors (30 vaccinated and 23 unvaccinated controls), two 68-1 RhCMV/SIV vaccine-protected RM used for $CD8^+$ cell depletion and two treatment-naïve RM used as adoptive transfer recipients. Fifteen of the unvaccinated negative controls and the 15 Rh67-intact 68-1 RhCMV/SIV vaccinated group used in the comparative immunogenicity/efficacy analysis were presented in detail in the companion manuscript (*13*) The 15 RM vaccinated ΔRh67 68-1 RhCMV vectors were vaccinated and SIV-challenged subsequent to the RM in the companion manuscript using the same and vaccination regimen and challenge protocol. To ensure comparability of the challenge procedure, these RM were challenged in parallel with an additional 8 unvaccinated controls. RhCMV vectors were routinely dosed at $10^6$-$10^7$ infectious units for immunogenicity analysis and $5x10^6$ infectious units per vector for large group immunogencity and efficacy analysis, all via subcutaneous administration. For the former studies the vaccine vector was given once; for latter studies, RM were subcutaneously inoculated with same vaccine vectors at the same dose a second time 18 weeks post-initial vaccination (homologous boost). SIV challenge and adoptive transfer analysis of replication competent SIV were performed as previous described (*8-10, 13*). Depletion of $CD8^+$ cells from RM was accomplished using treatment with the monoclonal antibody (mAb) M-T807R1 (10 mg $kg^{-1}$ subcutaneously at day 0, and 5 mg $kg^{-1}$ intravenously at days 3, 7, and 10), as previously described (*8, 9, 27*). At assignment, all study RM were free of cercopithicine herpesvirus 1, D-type simian retrovirus, simian T-lymphotrophic virus type 1, and Mycobacterium tuberculosis, and all but 1 (see **fig. S4**) were naturally RhCMV-infected. One RM, used to test immunogenicity of a Rh67-deleted 68-1 RhCMV vector also lacking viral inhibitors of MHC-Ia presentation (ΔRh178ΔRh182-9), which eliminates the ability to superinfect $RhCMV^+$ RM, was specially raised to be RhCMV-naïve at assignment (*9*). All study RM were housed at the Oregon National Primate Research Center (ONPRC) in Animal Biosafety level (ABSL)-2 (vaccine phase) and ABSL-2+ rooms (challenge phase) rooms with autonomously controlled temperature, humidity, and lighting. Study RM were both single and pair cage-housed. Animals were only paired with one another during the vaccine phase if they were from the same vaccination group. All RM were single cage-housed during the challenge phase due to the infectious nature of the study. Regardless of their pairing, all animals had visual, auditory and olfactory contact with other animals. Single cage-housed RM received an enhanced enrichment plan that was designed and overseen by NHP behavior specialists. RM were fed commercially prepared primate chow twice daily and received supplemental fresh fruit or vegetables daily. Fresh, potable water was provided via automatic water systems. Physical exams including body weight and complete blood counts were performed at all protocol time points. RM were sedated with ketamine HCl or Telazol for procedures, including intradermal and subcutaneous vaccine administration, venipuncture, bronchoalveolar lavage, bone marrow and lymph node biopsy and SIV challenge. At humane or scheduled endpoints, RM were euthanized with sodium pentobarbital overdose (>50 mg/kg) and exsanguinated via the distal aorta, and tissue collection at necropsy was performed by a certified veterinary pathologist. RM care and all experimental protocols and procedures were approved by the ONPRC Institutional Animal Care and Use Committee (IACUC). The ONPRC is a Category I facility. The Laboratory Animal Care and Use Program at the ONPRC is fully accredited by the American Association for Accreditation of Laboratory Animal Care (AAALAC) and has an approved Assurance (#A3304-01) for the care and use of animals on file with the NIH Office for Protection from Research Risks. The IACUC adheres to national guidelines established in the Animal Welfare Act (7 U.S.C. Sections 2131–2159) and the Guide for the Care and Use of Laboratory Animals (8th Edition) as mandated by the U.S. Public Health Service Policy.

**T cell assays.** To monitor CD8+ T cell responses to RhCMV-infected cells by flow cytometric intracellular cytokine staining (ICS) TRF were infected at an MOI of 3 for 24 hrs. Cells were harvested by trypsinization, resuspended at $4x10^6$ cells/ml in DMEM and 50μl aliquots of the cell suspensions

($2x10^5$ cells) were added to $5x10^5$ CD8β+ T cells in 50μl isolated from peripheral blood mononuclear cells (PBMC) of 68-1 RhCMV-immunized RM using a non-human primate $CD8^+$ T cell isolation kit (Miltenyi Biotec) and LS columns (Miltenyi Biotec) in the presence or absence of MHC blocking reagents.

SIV-specific $CD4^+$ and $CD8^+$ T cell responses were measured in PBMC or tissue-derived mononuclear cells by ICS as previously described (*11, 12*). Briefly, individual or whole protein mixes of sequential 15-mer peptides (11 amino acid overlap) spanning the $SIV_{mac239}$ Gag, 5'-Pol, Nef, Rev, Tat, and Vif proteins or individual $SIV_{mac239}$ Gag supertope peptides [$Gag_{211-222}$ (53), $Gag_{276-284}$ (69), $Gag_{290-301}$ (73), $Gag_{482-490}$ (120)] were used as antigens.

Mononuclear cells were incubated at 37°C with the peptide(s) or cells and co-stimulatory anti-CD28 (CD28.2, Purified 500 ng/test: eBioscience, Custom Bulk 7014-0289-M050) and anti-CD49d mAb (9F10, Purified 500 ng/test: eBioscience, Custom Bulk 7014-0499-M050) for 1 hr, followed by an additional 8h incubation in the presence of Brefeldin A (5 μg/ml; Sigma-Aldrich). Stimulation in the absence of peptides or cells served as background control. After incubation, stimulated cells were stored at 4°C until staining with combinations of fluorochrome-conjugated monoclonal antibodies including: anti-CD3 (SP34-2: Pacific Blue; BD Biosciences, Custom Bulk 624034 and PerCP-Cy5.5; BD Biosciences, Custom Bulk 624060), anti-CD4 (L200: FITC; BD Biosciences, Custom Bulk 624044 and AmCyan; BD Biosciences, Custom Bulk 658025), anti-CD8α (SK1: APC-Cy7; eBioscience, Custom Bulk 7047-0087-M002), anti-TNF-α (MAB11: APC; BD Biosciences, Custom Bulk 624076 and FITC; BD Biosciences, Custom Bulk 624046 and PE; BD Biosciences, Custom Bulk 624049), anti-IFN-γ (B27: APC; BD Biosciences, Custom Bulk 624078 and FITC; BD Biosciences, 554700) and anti-CD69 (FN50: PE; eBioscience, Custom Bulk CUST01282 and PE-TexasRed; BD Biosciences, Custom Bulk 624005) and for polycytokine analyses, anti-IL-2 (MQ1-17H12; PE Cy-7; BioLegend), and anti-MIP-1β (D21-1351, PBV421; BD Biosciences).

The MHC restriction (MHC-Ia, MHC-E, MHC-II) of $CD8^+$ T cell responses to peptides or RhCMV-infected cells was determined by pre-incubating isolated mononuclear cell aliquots or RhCMV-infected cells for 1 hr at room temperature (prior to adding peptides or combining effector and target cells and incubating per the standard ICS assay) in the presence (and absence) of each the following specific inhibitors: 1) the pan anti-MHC-I mAb W6/32 (10μg/ml), 2) the MHC-II-blocking mAb G46.6 (10μg/ml) or anti-HLA-DR (Clone L243, BioLegend #92203, 1mg/ml)) and 2μl CLIP peptide (MHC-II-associated invariant chain, amino acids 89-100, Intavis, 1mg/ml) per test sample, or 3) the MHC-E blocking VL9 peptide (VMAPRTLLL; 20μM). Stimulated cells were fixed, permeabilized, stained and analyzed as described above. To be considered MHC-E-restricted by blocking, the individual peptide response must have been blocked by both anti-pan MHC-I clone W6/32 and MHC-E-binding peptide VL9, and not blocked by anti-MHC-II. MHC-II-restricted responses were blocked by anti-MHC-II but not anti-MHC-I or VL9, and MHC-Ia-restricted responses were blocked by anti-MHC-Ia only (*11, 12*). Responses that did not meet these inhibition criteria were considered indeterminate.

For analysis of memory differentiation (central- vs transitional- vs effector-memory) of SIV Gag-specific $CD4^+$ and $CD8^+$ T cells, PBMC were stimulated with SIV Gag 15mer peptide mix as described above, except that the CD28 co-stimulatory mAb was used as a fluorochrome conjugate to allow CD28 expression levels to be later assessed by flow cytometry, and in these experiments, cells were surface-stained after incubation for lineage markers CD3, CD4, CD8, CD95 and CCR7 (see below for mAb clones) prior to fixation/permeabilization and then intracellular staining for response markers (CD69, IFN-γ, TNF-α; note that Brefeldin A treatment preserves the pre-stimulation cell-surface expression phenotype of phenotypic markers examined in this study).

Stained samples were analyzed on an LSR-II flow cytometer (BD Biosciences). Data analysis was performed using FlowJo software (Tree Star). In all analyses, gating on the lymphocyte population was followed by the separation of the $CD3^+$ T cell subset and progressive gating on $CD4^+$ and $CD8^+$ T cell subsets. Antigen-responding cells in both $CD4^+$ and $CD8^+$ T cell populations were determined by their intracellular expression of CD69 and either or both of the cytokines IFN-γ and TNF-α (or in polycytokine analyses, expression of CD69 and any combination of the cytokines: IFN-γ, TNF-α, IL-2, MIP-1β). After background subtraction, the raw response frequencies above the assay limit of detection were "memory-corrected" (e.g., % responding out of the memory population), as previously described (*8-10, 38*), using combinations of the following fluorochrome-conjugated mAbs to define the memory vs naïve subsets CD3 (SP34-2; Alexa700, PerCP-Cy5.5), CD4 (L200; AmCyan), CD8α (SK-1; APC, PerCP-cy-5.5), TNF-α (MAB11; FITC), IFN-γ (B27; APC), CD69 (FN50; PE), CD28 (CD28.2; PE-TexasRed), CD95 (DX2; PE), CCR7 (15053; Pacific Blue), and Ki67 (B56; FITC). For memory phenotype analysis of SIV Gag-specific T cells, all $CD4^+$ or $CD8^+$ T cells expressing CD69 plus IFN-γ and/or TNF-α were first Boolean OR gated, and then this overall Ag-responding population was subdivided into the memory subsets of interest on the basis of surface phenotype (CCR7 vs CD28). Similarly, for polycytokine analysis of SIV Gag-specific T cells, all $CD4^+$ or $CD8^+$ T cells expressing CD69 plus cytokines were Boolean OR gated and polyfunctionality was delineated with any combination of the four cytokines tested (IFN-γ, TNF-α, IL-2, MIP-1β) using the Boolean AND function.

**SIV detection assays.** Plasma SIV RNA levels were determined using an SIV Gag-targeted quantitative real time/digital RT-PCR format assay, essentially as previously described, with 6 replicate reactions analyzed per extracted sample for assay thresholds of 15 SIV RNA copies/ml (*9, 10, 39*).

**Statistical Analysis**. Boxplots show jittered points and a box from 1st to 3rd quartiles (IQR) and a line at the median, with whiskers extending to the farthest data point within 1.5*IQR above and below the box. For all comparisons of T cell response parameters, we performed Wilcoxon rank-sum tests comparing the ΔRh67 68-1 RhCMV/SIV vector vaccinated group to the Rh67-intact 68.1 RhCMV/SIV vector vaccinated reference group. For longitudinal responses, we calculated the area under the curve (AUC) or the plateau average value for each RM, as denoted in the figure legends. All Wilcoxon P values are based on two-sided tests and were adjusted using the Holm procedure for family-wise error rate control. P values for analyses of efficacy were based on two-sided exact tests of binomial proportions. Analyses were performed in R v3.6.0 with the package Exact v2.0 (*40*).

### SUPPLEMENTAL FIGURES

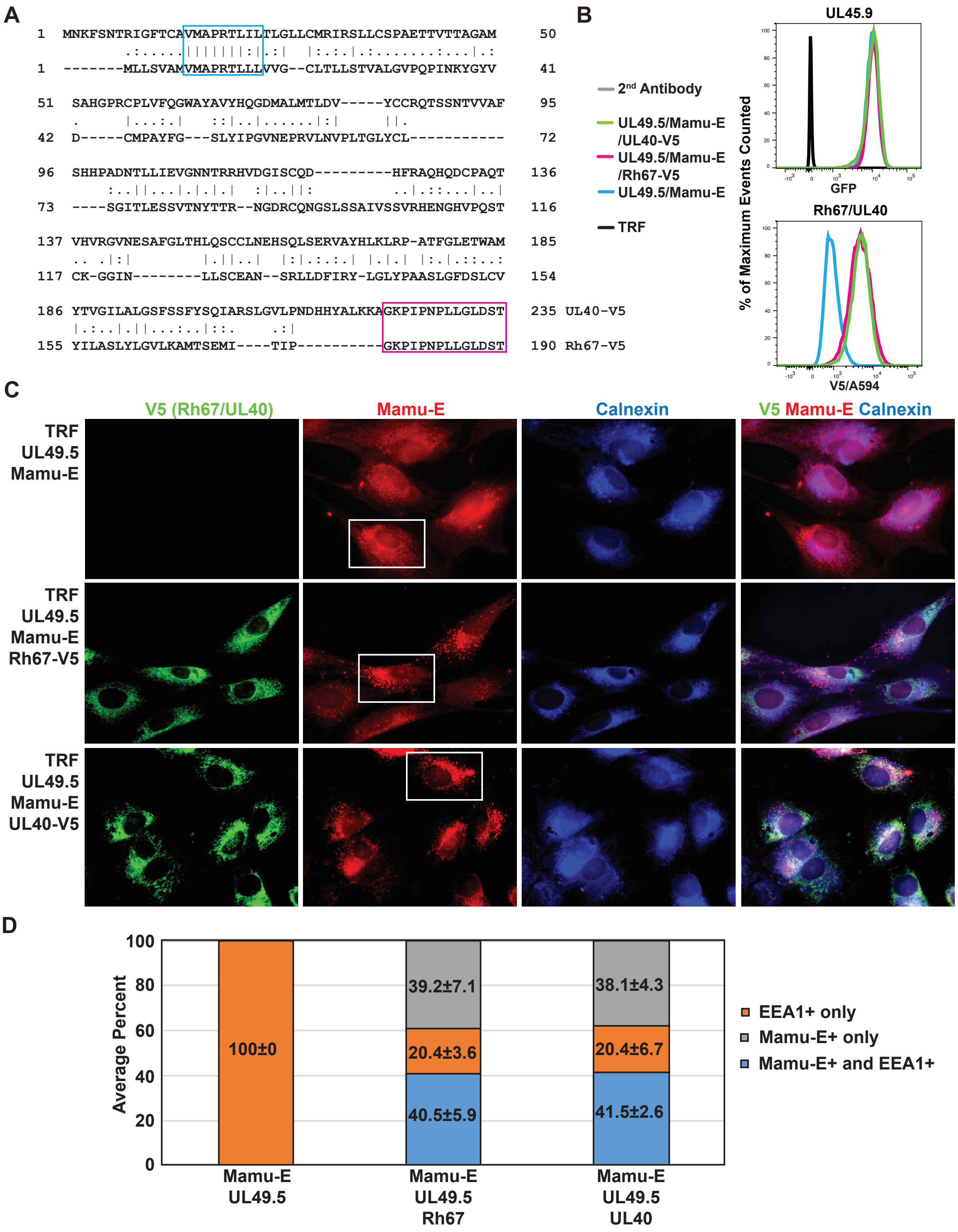


**Figure S1: Rh67 and UL40 contain a conserved VL9 sequence and cause TAP-independent Mamu-E transport and internalization. A)** V5-tagged Rh67 (Accession #AFL03561.1) and UL40 (Accession #QHB20491.1) were aligned using the global alignment with free end gap tool (Geneious). The embedded VL9 sequence and added V5-epitope tags are highlighted. **B)** Flow cytometric analysis of

Rh67 and UL40-transduced cells for expression of GFP, which is co-expressed with UL49.5, and for the V5 tag. Surface expression of Mamu-E (**Fig. 1C**) was determined on $GFP^{+}$ and $V5^{+}$ cells. **C)** Intracellular localization of V5-taggegd Rh67 and UL40, Calnexin and Mamu-E was determined in UL49.5-expressing, Mamu-E-transfected TRF by immunofluorescence assay using indicated antibodies. Nuclei were stained with DAPI. Cells shown in **Fig. 1D** are highlighted. **D)** Vesicular staining of MHC-E and EEA1 upon internalization of MHC-E/antibody complexes (**Fig. 1E**) was quantified by counting all EEA1 and/or Mamu-E positive vesicles in four individual cells from three independent experiments for each cell line. Average percentages of vesicles that are $EEA1^{+}$ only, Mamu-$E^{+}$ only, and dual $EEA1^{+}$ and Mamu-$E^{+}$ are shown (+/- SEM) for each cell line.

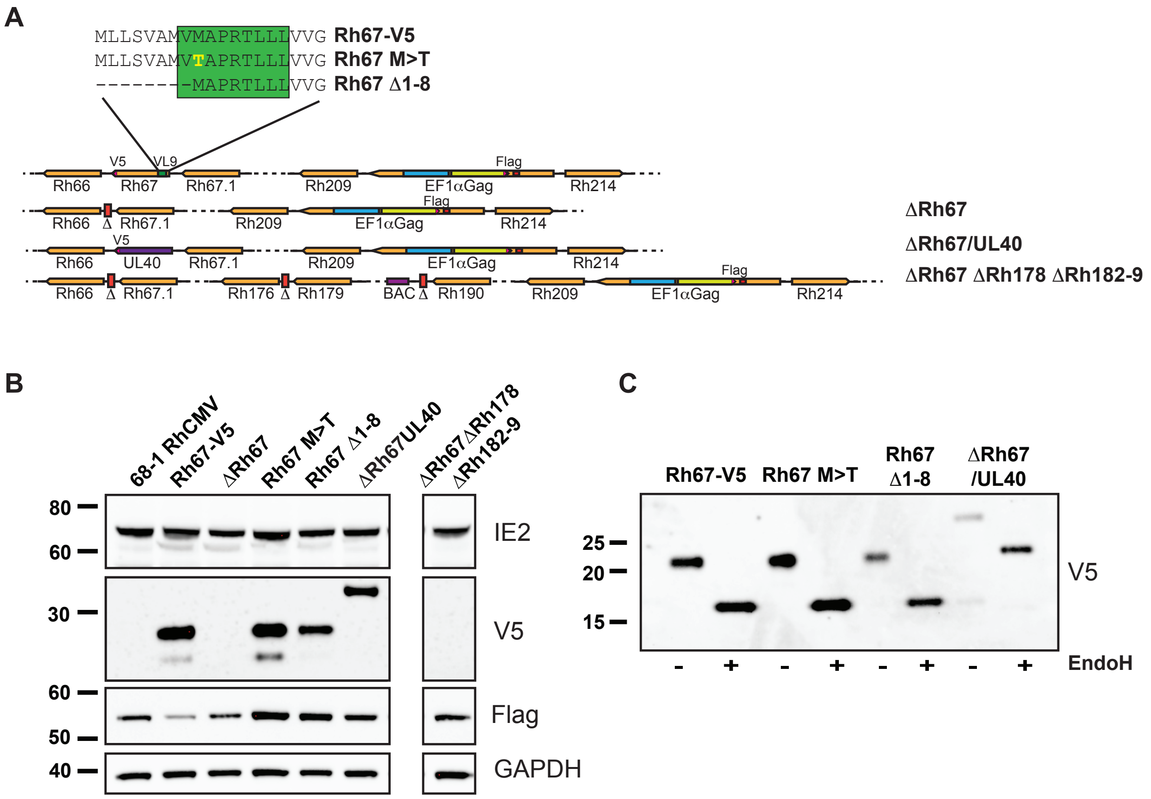


**Figure S2: Description and characterization of Rh67-modified 68-1 RhCMV/SIVgag recombinants. (A)** Schematic of RhCMV recombinants. Rh67-V5 (GenBank: MN622886) was generated by replacing Rh67 with Rh67 containing a C-terminal V5 epitope tag in 68-1 RhCMV/SIVgag (Genbank: MN437483) which contains EF1α-driven, FLAG-tagged SIVgag in Rh211 as previously described (*5*). For Rh67M>T (MN622885), Rh67Δ1-8 (MN622887) or ΔRh67/UL40 (MN622884) the Rh67 gene in 68-1 RhCMV/SIVgag was similarly replaced with V5 epitope-tagged Rh67 variants or UL40 whereas Rh67 was deleted from the same vector to generate ΔRh67 (MN622881). Note that the N-terminal methionine in Rh67Δ1-8 will be removed co-translationally so that both V and M are missing from the mature protein. ΔRh67ΔRh178ΔRh182-9 (MT068444) was also derived from 68-1 RhCMV/SIVgag by deleting Rh67, Rh178 and Rh182-189. (**B)** Immunoblot of TRF infected with the indicated constructs (MOI=3) for 48 hrs prior to lysis, SDS-PAGE and immunoblot using the indicated antibodies specific for RhCMV IE, V5-tagged Rh67 and UL40 proteins, FLAG-tagged SIVgag and cellular GAPDH. (**C)** TRF were infected and harvested as in B). Where indicated, lysates were treated with EndoH prior to electrophoretic separation and immunoblot with anti-V5 antibody.

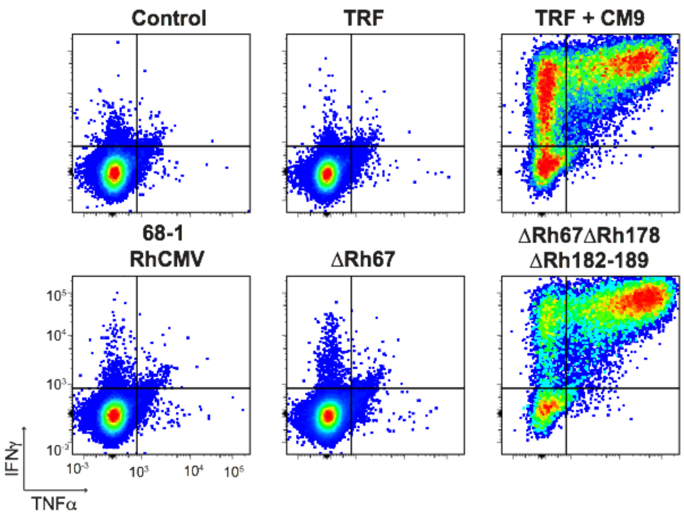


**Figure S3: Triggering of Mamu-A01-restricted, SIV Gag CM9-specific, CD8$^+$ T cells by RhCMV recombinants.** A CD8$^+$ T cell line specific for the Mamu-A*01-restricted Gag CM9 epitope was generated as described (*41*), and co-cultured with Mamu-A*01$^+$ telomerized rhesus fibroblasts (TRF) pulsed with 20nM CM9 peptide for 90 min or infected with the indicated SIVgag-expressing 68-1 RhCMV recombinants for 24 hours prior to ICS. Dot plots are gated on CD3$^+$ CD8$^+$ T cells following co-culture with the indicated cells.

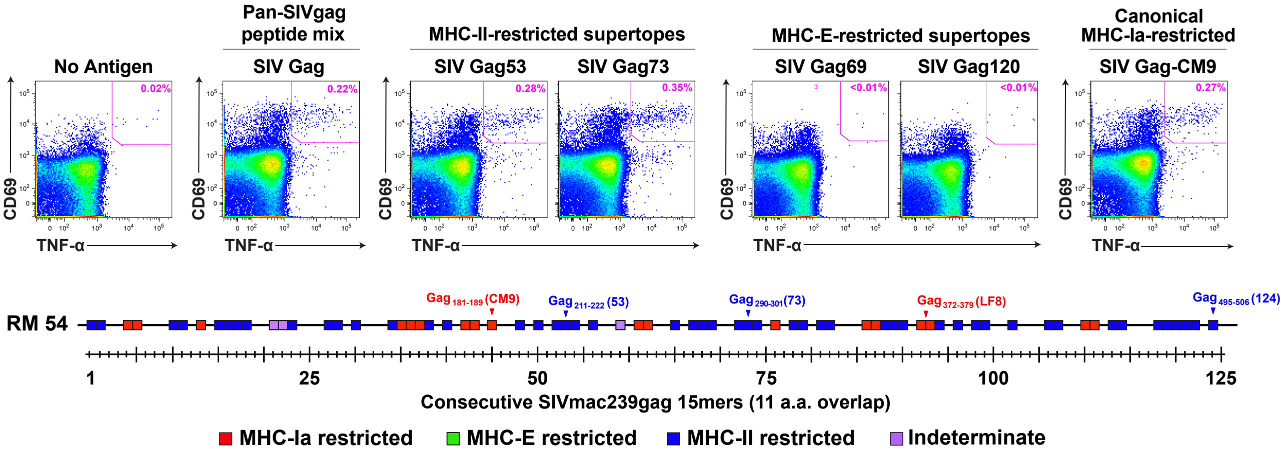


**Figure S4. Requirement for Rh67 for MHC-E-restricted $CD8^+$ T cell priming in the absence of all viral inhibitors of MHC-Ia presentation.** Analysis of SIV Gag-specific $CD8^+$ T cell responses elicited by a ΔRh67ΔRh178ΔRh182-9 68-1 RhCMV/SIVgag vector in a *Mamu* A*01$^+$, RhCMV-naïve RM [Note: RhCMV-naïve RM required because RhCMV vectors lacking all viral inhibitors of MHC-Ia presentation do not super-infect RhCMV$^+$ RM, as previously described (*9*)]. Peripheral blood mononuclear cells from this RM were assessed by a flow cytometric intracellular cytokine staining for response to 1) a mixture of 125 consecutive 15mer peptides comprising the SIV Gag protein sequence (pan-SIVgag peptide mix), 2) individual MHC-E and MHC-II-restricted SIV Gag supertopes, 3) an immunodominant, canonical *Mamu* A*01-restricted epitope ($Gag_{181-189}$ CM9) (top panels) and 4) each of 125 consecutive, overlapping 15mer SIV Gag peptides (bottom panel). In the top panels, the flow cytometric profiles of CD69 vs. TNF-α are shown, with CD69$^+$, TNF-α$^+$ Ag-responding cells delineated by the pink boxes (%+ shown). In the bottom panel, above threshold responses to each of the individual consecutive 15mer peptides ($\geq$ 0.05% after background subtraction) are indicated by a box, which is colored green, blue or red based on blocking of the response by the MHC-E blocking peptide VL9, the anti-MHC-II mAb G46-6, and/or the anti-MHC-Ia mAb W6/32. The positions of the MHC-II-restricted supertopes and two known *Mamu* A*01-restricted canonical SIV Gag epitopes are indicated in the figure.

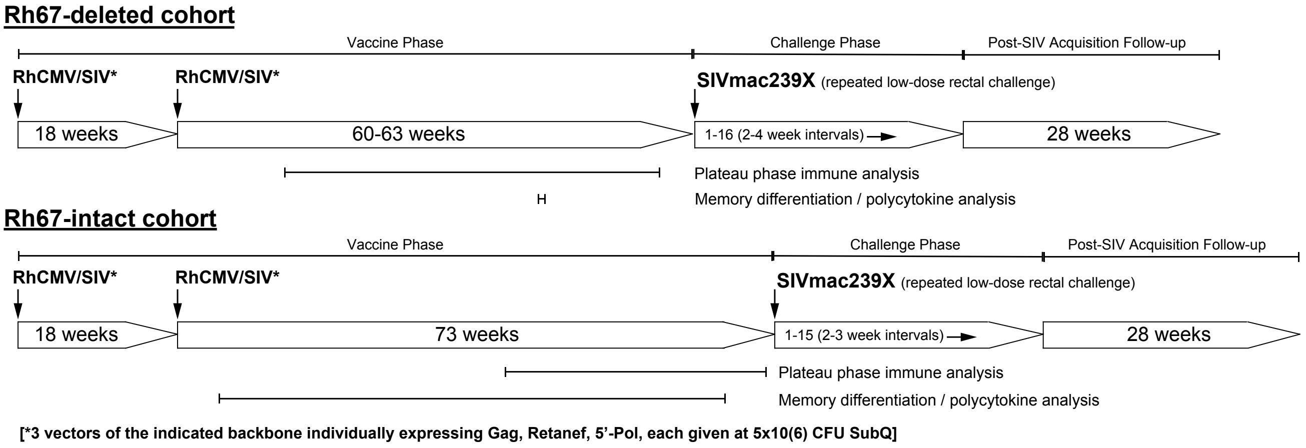


**Figure S5. Protocols for the comparison of the immunogenicity and efficacy of Rh67-deleted vs. Rh67-intact RhCMV/SIV vectors.** The positive control group for the efficacy assessment of ΔRh67 RhCMV/SIV vectors is the 68-1 RhCMV/SIV vaccinated cohort reported in the companion manuscript (*13*). This study was performed separately from the evaluation of the Rh67-deleted vaccine; however, key aspects of these studies were essentially identical, including vaccine inserts, vaccine dose, vaccine boost, challenge stock, challenge frequency and challenge monitoring. Primary differences include a slightly shorter vaccine phase for the ΔRh67 vector vaccinated cohort, and the performance of specialized immune assays on different timepoints during plateau phase. Of note, both vaccine cohorts were challenged with their own separate concurrent unvaccinated controls: n=15 for the Rh67-intact vaccine; n=8 for the ΔRh67 vaccine.

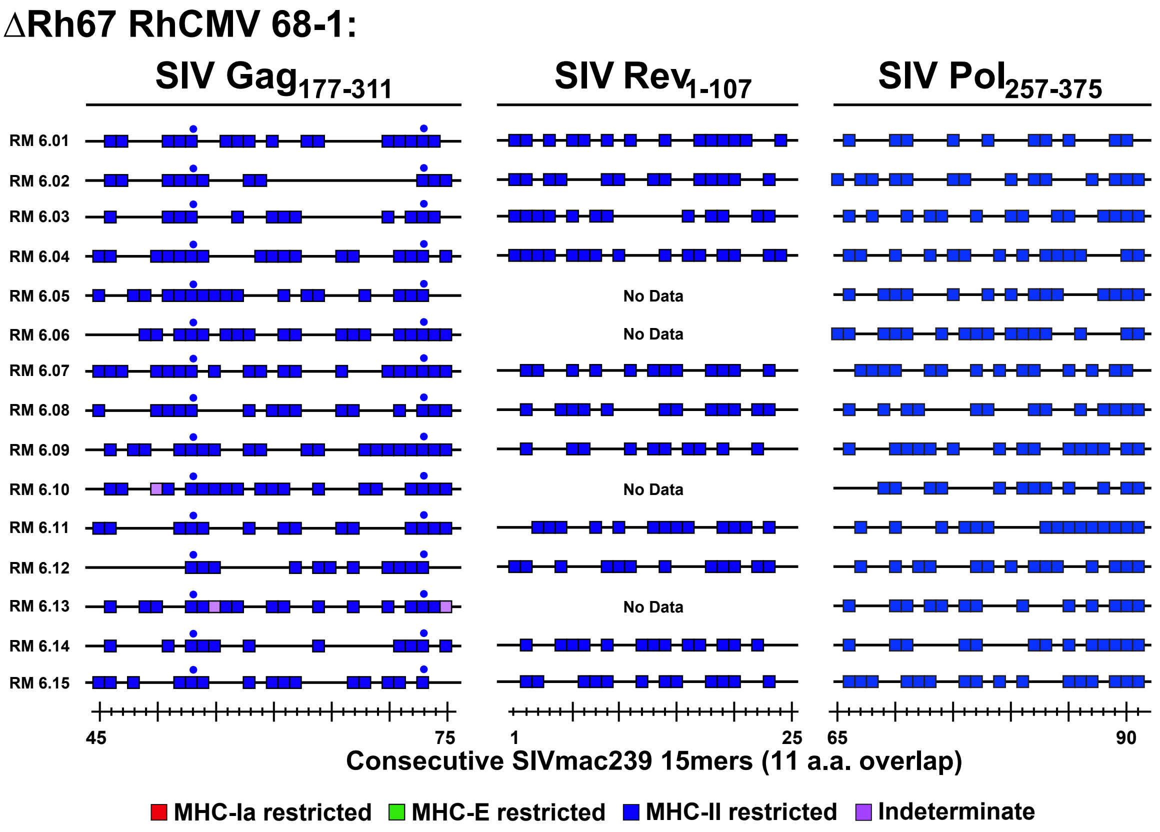


**Figure S6. Epitope analysis of ΔRh67 68-1 RhCMV/SIV vaccine-elicited, SIV-specific CD8$^+$ T cell responses in RMs destined for SIV challenge.** The epitope targeting characteristics of the CD8$^+$ T cell responses elicited by ΔRh67 68-1 RhCMV/SIV in the RM destined for SIV challenge were analyzed for indicated portions of each insert (SIV Gag$_{177\text{-}311}$, SIV Rev$_{1\text{-}107}$, SIVp Pol$_{257\text{-}375}$ for the SIVgag, SIVretanef, and SIV5'-pol inserts, respectively, corresponding to 15mers 45-75, 1-25, and 65-91). As indicated by the exclusively blue boxes, blocking analysis (see Methods) indicated that all epitope-specific responses to the SIV inserts in all 3 vectors used in each RM were presented by MHC-II. The blue dots indicate that MHC-II-restricted supertope responses were independently confirmed with optimal peptides.

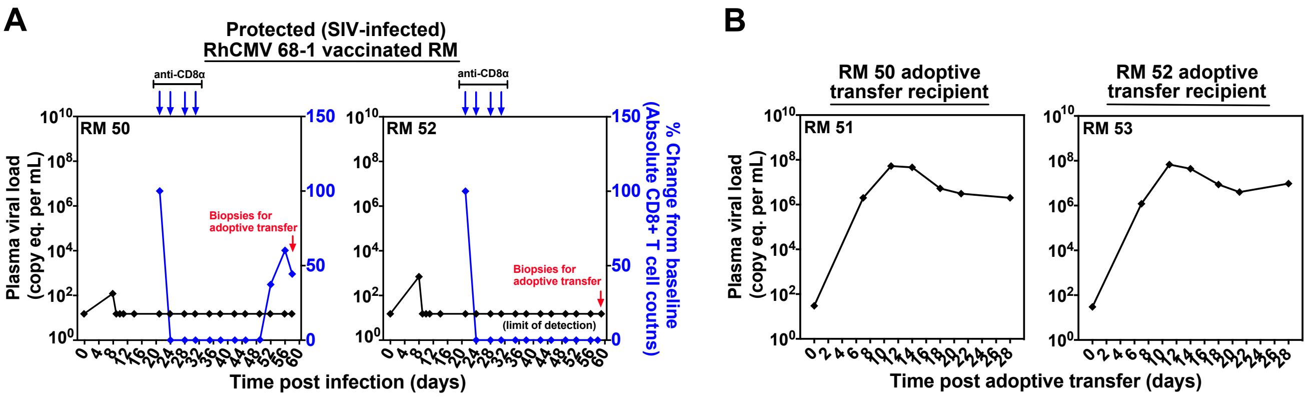


**Figure S7. Resistance of 68-1 RhCMV/SIV vaccine-induced replication arrest to *in vivo* CD8α⁺ cell depletion.** (**A**) Black lines delineate plasma viral load profiles (pvl; limit of detection = 30 copies/ml) of two 68-1 RhCMV/SIV vector vaccinated RM (RM 50 and RM 52) that were protected following productive SIV challenge by validated criteria (pvl blip at day 8 and development of SIVvif-specific T cell responses in the absence of sustained plasma viremia in both RM) (*8-10*). Both protected RM were treated with the CD8α mAb M-T807R1 while aviremic (pvl below limit of detection) at day 21 (10mg/kg), 24 (5mg/kg), 28 (5mg/kg), and 31 (5mg/kg), effectively depleting circulating $CD8^+$ T cells for nearly 4 weeks in RM 50 and through the end of observation in RM 52. No subsequent plasma viremia was detected during/after CD8α cell depletion. (**B**) To confirm that these RM were SIV-infected and in RhCMV/SIV vaccine-induced "replication arrest" during depletion, peripheral lymph node and bone marrow biopsies were obtained at day 60 post-infection from both RM for adoptive transfer analysis of arrested SIV infection (*10*). The figure shows pvl profiles of two SIV-naïve recipient RM, RM 51 and RM 53, after intravenous administration of 60 million cells (30 million bone marrow cells and 30 million lymph node cells) obtained from the biopsies of RM 50 and RM 52, respectively. Both recipient RM 51 and RM 53 developed typical progressive SIV infection after this cellular transfer, indicating the presence of replication competent SIV in the protected donor RMs throughout the period of $CD8\alpha^+$ cell depletion.
